# Here be dragons: important spatial uncertainty driven by climate data in forecasted distribution of an endangered insular reptile

**DOI:** 10.1101/2021.06.14.448338

**Authors:** Nicolas Dubos, Stephane Augros, Gregory Deso, Jean-Michel Probst, Jean-Cyrille Notter, Markus A. Roesch

## Abstract

The effect of future climate change is poorly documented in the tropics, especially in mountainous areas. Yet, species living in these environments are predicted to be strongly affected. Newly available high-resolution environmental data and statistical methods enable the development of forecasting models. Nevertheless, the uncertainty related to climate models can be strong, which can lead to ineffective conservation actions. Predicted studies aimed at providing conservation guidelines often account for a range of future climate predictions (climate scenarios and global circulation models). However, very few studies considered potential differences related to baseline climate data and/or did not account for spatial information (overlap) in uncertainty assessments. We modelled the environmental suitability for *Phelsuma borbonica*, an endangered reptile native to Reunion Island. Using two metrics of species range change (difference in overall suitability and spatial overlap), we quantified the uncertainty related to the modelling technique (n = 10), sample bias correction, climate change scenario, global circulation models (GCM) and baseline climate (CHELSA *versus* Worldclim). Uncertainty was mainly driven by GCMs when considering overall suitability, while for spatial overlap the uncertainty related to baseline climate became more important than that of GCMs. The uncertainty driven by sample bias correction and variable selection was much higher when assessed based on spatial overlap. The modelling technique was a strong driver of uncertainty in both cases. We eventually provide a consensus ensemble prediction map of the environmental suitability of *P. borbonica* to identify the areas predicted to be the most suitable in the future with the highest certainty. Predictive studies aimed at identifying priority areas for conservation in the face of climate change need to account for a wide panel of modelling techniques, GCMs and baseline climate data. We recommend the use of multiple approaches, including spatial overlap, when assessing uncertainty in species distribution models.

## Introduction

Predictive studies of climate change effects are poorly documented in the tropics, especially in rare and endangered species (Pearson et al., 2014). Yet, tropical species are predicted to be more severely impacted by climate change than temperate species because they live closer to their thermal limit (Tewksbury, Huey, & Deutsch, 2008; Jiguet et al., 2010; Dubos et al., 2019). This is particularly true for species with a narrow distribution, a narrow niche, or living in highly heterogeneous environments such as mountainous areas (Raxworthy et al., 2008; Freeman & Class Freeman, 2014; Platts et al., 2014; Tang et al., 2018; Ahmadi et al., 2019; Cordier et al., 2020; Manes et al., 2021). Those species are difficult to model mostly because of data scarcity and statistical constrains related to low sample sizes and the lack of spatial coverage in environmental gradients. The recent availability of high-resolution environmental data along with methodological advances help filling this gap of knowledge (e.g., Dubos et al., 2021; Lannuzel et al., 2021). Forecasting models are useful to anticipate conservation management in the face of climate change. Specifically, species distribution models (SDMs) can be used to identify priority areas for protection (Leroy et al., 2014; Lannuzel et al., 2021), suitable conditions for habitat restoration and species (re)introductions and translocations (Minteer & Collins, 2010; Wilson, Roberts, & Reid, 2011; Adhikari, Barik, & Upadhaya, 2012; Draper, Marques, & Iriondo, 2019; Bellis et al., 2020; Butt et al., 2020; Westwood et al., 2020; Zhong et al., 2021).

A range of climate data is available for species distribution models, including several baseline climate data, some of which are used to predict climate change (e.g., Worldclim and CHELSA; Fick & Hijmans, 2017; Karger et al., 2017). Future climate predictions include multiple emission gas scenarios (Shared Socioeconomic Pathways, SPP; as known as Representative Concentration Pathways, RCP) and global circulation models (GCMs). Both gas emission scenarios and GCMs can produce highly heterogeneous results in terms of predicted future distributions (Buisson et al., 2010; Baker et al., 2016). Most predictive studies included a range of scenarios and GCMs, but very few have considered potential uncertainties related to baseline climate data (Baker et al., 2016; Morales-Barbero & Vega-Álvarez, 2019; Datta, Schweiger, & Kühn, 2020; Ocon, 2020; Dubos, et al., 2021). This can induce a risk of misidentification of suitable environments, which can affect conservation prioritisation (Kujala et al., 2013; Baker et al., 2016; Muscatello, Elith, & Kujala, 2020) and subsequently lead to ineffective conservation actions (Converse & Sipe, 2021). Despite recent studies emphasising the need to account for multiple baseline climates (Morales-Barbero & Vega-Álvarez, 2019; Datta, Schweiger, & Kühn, 2020; Ocon, 2020), most studies aimed at providing conservation guidelines do not assess the uncertainty related to bioclimatic input data.

Uncertainty assessments in SDMs can be made by comparing a range of modalities in model settings or data. These comparisons can be based on differences in performance metrics such as the area under the receiving operating characteristic curve (AUC) or the true skill statistic (TSS; Tessarolo et al., 2021). However, such performance comparisons require independent data to be reliable. In absence of independent data, uncertainty can be assessed based on differences in predicted species range change between model input parameters, climate scenarios or climate data (e.g., Kujala et al., 2013; Baker et al., 2016; Muscatello, Elith, & Kujala, 2020). Species range change can be estimated using the difference between current and future summed suitability. This approach enables to identify which parameter drives the highest uncertainty in terms of overall suitability across a region. However, it ignores the spatial component in suitability differences. For instance, a difference in shift in distribution with no change in overall suitability would remain undetected. Another approach is to discount the variation between model replicates from mean suitability scores (Kujala et al., 2013). This method is highly relevant for conservation purposes since it provides a spatially explicit map of the most consistently identified suitable areas but prevents from disentangling the sources of uncertainty. A possible approach to assess each source of uncertainty while accounting for spatial information is the use of overlap metrics such as Pearson’s coefficient or similarity indices (Muscatello, Elith, & Kujala, 2020; Dubos, Montfort, et al., 2021). To date, no study has tested for potential differences in uncertainty assessments between these approaches.

In our study we generate a set of SDMs for the two subspecies of the Reunion Island day gecko, *Phelsuma borbonica borbonica* and *P. borbonica mater* (subsequently referred to as *P. borbonica*), and test potential differences in uncertainty assessments between the different approaches. Reunion Island is located in the western Indian Ocean and has faced a number of alterations over the last century related to agricultural practices (mostly sugarcane cultivation) and invasive species (Strasberg et al., 2005; Dubos, 2013; Fenouillas et al., 2021; Irl et al., 2021). Its herpetofauna has already been strongly modified after multiple local extinctions events and the arrival of allochthonous species (Cheke, 1987; Cheke & Hume, 2010; Sanchez & Probst, 2016). Today only two native reptile species remain, the indigene *P. brobonica* and the endemic *P. inexpectata*, while five reptile species having faced extinction in the recent past (Cheke & Hume, 2010). Climate change can facilitate biological invasions and alters habitat suitability for locally adapted species (Mainka & Howard, 2010; Gillard et al., 2017). To date, very few studies have focused on the potential effects of climate change on the future of Reunion Island’s biodiversity (but see Legrand et al., 2016). Dubos et al. (2021) predicted the potential extinction of the endemic Reunion Island reptile, *P. inexpectata*, driven by climate change.

The Reunion Island day gecko, *P. borbonica* is classified nationally as endangered due to its narrow distribution. With high rates of land use change in Reunion Island, fire hazards and high pressure related to introduced species (Macdonald & Cedex, 1991; Strasberg et al., 2005; Lagabrielle et al., 2009; Sanchez & Probst, 2016), many populations are isolated and are exposed to local extinctions. This calls the urgent need to identify priority areas for habitat conservation and restoration.

We assess for the first time the climatic niche of *P. borbonica* and the potential effects of climate change on its future distribution. We quantify the uncertainty in species range change (SRC) driven by the statistical methods (modelling technique, sample bias correction, variable selection) and climate data (RCPs, GCMs and baseline data) using two approaches (difference in summed suitability and spatial overlap) in the predicted future distribution of *P. borbonica*. We eventually provide a consensual map accounting for uncertainty to guide conservation actions.

## Methods

### Occurrence data

*Phelsuma borbonica* is mostly distributed in the forested areas of the eastern, southern, and northern parts of Reunion Island, from sea level to 2800 m (Meier, 1995; Mickaёl Sanchez & Probst, 2017). Most observations are made in intermediate altitudes, near trails, on artificial structures or at the edge of pristine or disturbed forested areas (Augros et al., 2017). We obtained 5922 occurrence data (2648 after removing redundancies) from the Système d’information de l’inventaire du patrimoine naturel (SINP) provided by the Direction de l’Environnement, de l’Aménagement et du Logement (DEAL) and the Parc National de la Réunion (PNRun) and some additional unpublished occurrence data from various sources (Biotope Océan Indien, Cynorkis, Eco-Med Océan Indien, Nature Océan Indien). We removed 171 observations corresponding to single observations with no evidence of persistence (presumably corresponding to individuals transported out of their native range; Deso, 2001), translocated individuals, inaccurate coordinates, or indices of past presence (e.g., subfossil clutches). To limit spatial autocorrelation, we resampled one occurrence per occupied pixel at the resolution of the environmental data (3 arc seconds, approximately 100m). The final sample included 1050 points when sampled from the Worldclim data, 1056 with CHELSA.

### Land use data

We accounted for the habitat requirements of our model species by applying a filter to the climate data. We obtained very high-resolution land cover categories (Urban, agricultural, natural, water) at 1.5m resolution (resampled at 100m for computing purposes) derived from remote sensing (Dupuy, Gaetano, & Le Mézo, 2020; Fig. Sx). We removed agricultural and urbanised areas, because *P. borbonica* is only found in natural and semi natural habitats, which enabled to minimise model complexity while remaining biologically realistic and relevant for conservation applications.

### Climate data

We used 19 bioclimatic variables for 30 arc sec (approximately 900m) resolution of the current climate data and of the 2070 projections from CHELSA (Karger et al., 2017) and from Worldclim global climate data (Fick & Hijmans, 2017). We removed isothermality (bio2) from the analysis because of a lack of variability in Reunion Island. We decided to include all the remaining variables because both temperature and precipitation are related to the species’ biology, including those referring to indices of variability. We used three Global Circulation Models (GCMs; i.e., BCC-CSM1-1, MIROC5 and HadGEM2-AO) and two greenhouse gas emission scenarios (the most optimistic RCP26 and the most pessimistic RCP85; Fig. S3, S4). To enable the filtering of unsuitable habitats, we downscaled the climate data to fit the resolution of the land cover variables (3 arc sec) using bilinear interpolation.

### Distribution modelling

We modelled and projected species distributions in R (version 4.0.3; R Core Team, 2020) with the Biomod2 R package (Thuiller et al., 2009), using 10 modelling techniques: generalised linear and generalised additive models (GLM and GAM; Guisan, Edwards, & Hastie, 2002), classification tree analysis (CTA; Prasad, Iverson, & Liaw, 2006), artificial neural network (ANN; Manel, Dias, & Ormerod, 1999), surface range envelop (SRE, also known as BIOCLIM; Booth et al., 2014), flexible discriminant analysis (FDA; Manel, Dias, & Ormerod, 1999), random forest (RF; Prasad, Iverson, & Liaw, 2006), multiple adaptive regression splines (MARS; Leathwick et al., 2005), generalised boosting model (GBM; Elith, Leathwick, & Hastie, 2008) and maximum entropy (MaxEnt; Phillips & Schapire, 2006). We generated five different sets of 1050/1056 pseudo-absences (to equal the number of presences) and ran a first set of models using a random pseudo-absence selection (Wisz & Guisan, 2009).

To accounting for sample bias, we reperformed all calculations using a non-random pseudo-absence selection. Following Phillips et al. (2009), we produced five sets of pseudo-absences selected around the presence points to reproduce the spatial bias of the sample. We used a geographic null model generated with the dismo R package (Hijmans, 2012) and used it as a probability weight for pseudo-absence selection. Since no independent data is available to assess the effect of sample bias correction, we used the relative overlap index (ROI; Dubos, Préau, et al., 2021) based on Schoener’s D overlap. The ROI enables to assess whether the effect of correction is significant compared to the variability between model runs. It computes (1) the mean overlap between the uncorrected and the corrected predictions (i.e., the absolute effect of correction), and (2) the overlap between every pair of model replicate (between each pseudo-absence and cross validation runs, individually for each modelling technique, i.e. model stochasticity). We computed the ROI as follows:

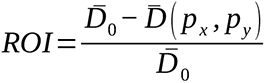

Where 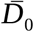 is the mean overlap between model runs of the corrected group and 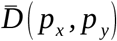 is the mean overlap between runs of the uncorrected and the corrected models. A value close to 0 represents a perfect match between predictions (i.e. no effect of sample bias correction). The overlaps between uncorrected and corrected groups tend to be significantly smaller than the overlaps between runs when the ROI approaches 1 (i.e. strong effect of sample bias correction).

For each individual baseline climate data, we selected one variable per group of inter-correlated variables to avoid collinearity (Pearson’s r > 0.7, Dormann et al., 2013) and assessed the relative importance of each variable kept with 10 permutations per model replicate (total = 500 for each baseline data). The variables included in the final models were those with a relative importance > 0.2 across at least 50% of model runs. To determine whether variation in future predictions was driven by baseline climate *per se* or by differences in the selected variables, we swapped the selected variables between baseline climates and repeated the whole process.

Model evaluation-We spatially partitioned the data into five folds, with three runs of block cross-validation (i.e., k-fold cross-validation; Fig. S1). We assessed model performance using the Boyce index, assumed to be the best evaluation metric with pseudo-absence data (Leroy et al., 2018). A value of 1 means the models reliably predicted the presence points while a value of 0 means that models did not perform better than random. For ensemble models, we excluded models for which the Boyce index was below 0.5 (Gillard et al., 2017). We verified that models are well informed for predictions on novel (future) data using clamping masks.

Uncertainty analysis–We assessed the uncertainty in species range change (SRC) related to the modelling technique, sample bias correction, climate scenarios, GCMs, baseline climate and variable selection. We quantified SRC following two approaches. Firstly, we computed SRC as the difference between the summed suitability scores of the baseline climate and the future predictions. Then we used linear models (LM, assuming Gaussian errors), with SRC as a response variable, and the aforementioned sources of uncertainty as explanatory variables, following Baker et al. (2016). We then assessed the proportion of deviance explained by each source of uncertainty *f* as follows:

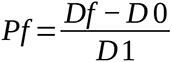

where, *Pf* = proportion of deviance explained by factor *f, D1* = deviance of full model, *Df* = deviance of full model minus factor *f* and *D0* = deviance of null model.

Secondly, we quantified SRC using an overlap metric. We computed the Schoener’s D overlap between baseline projections and future projections using the ENMTools R package (Warren, Glor, & Turelli, 2010). We eventually repeated the assessment of the proportion of deviance explained by the five aforementioned sources of uncertainty using beta-regression GLM instead of LM, since overlap measures range continuously between 0 and 1 (glmmTMB R package; Brooks et al., 2019).

In total, we computed 2000 models (10 modelling techniques * 5 pseudo-absence runs * 5 block crossvalidation runs * 2 modalities of sample bias correction * 2 baseline climate data * 2 modalities for variable selection) for the current distribution, and 12000 projections on future climate data (2000 models * 3 GCMs * 2 emission scenarios). To provide the most certain conservation guidelines, we built a consensus map of the mean predictions across all simulations (of the corrected group and with the original selected variables) after removing poorly performing models, discounted with the standard deviation (Kujala et al., 2013).

## Results

### Current distribution

Models predicted well the presence of *P. borbonica* (median Boyce index (Worldclim) = 0.81; median Boyce index (CHELSA) = 0.78; Fig. S2, S3). A few points, corresponding to isolated populations, fell into areas predicted as moderately suitable. Sample bias correction slightly increased the predictions of suitable areas, but its effect on projections was not significant compared to within-model variation (ROI = 0.08; Fig. S4). With the Worldclim baseline climate, the selected variables were mean annual temperature (Bio1) and winter precipitation (Bio19; Fig. 1). With the CHELSA baseline climate, the selected variables were mean annual temperature (Bio1) and summer precipitation (Bio18; Fig 2).

**Fig. 1.**
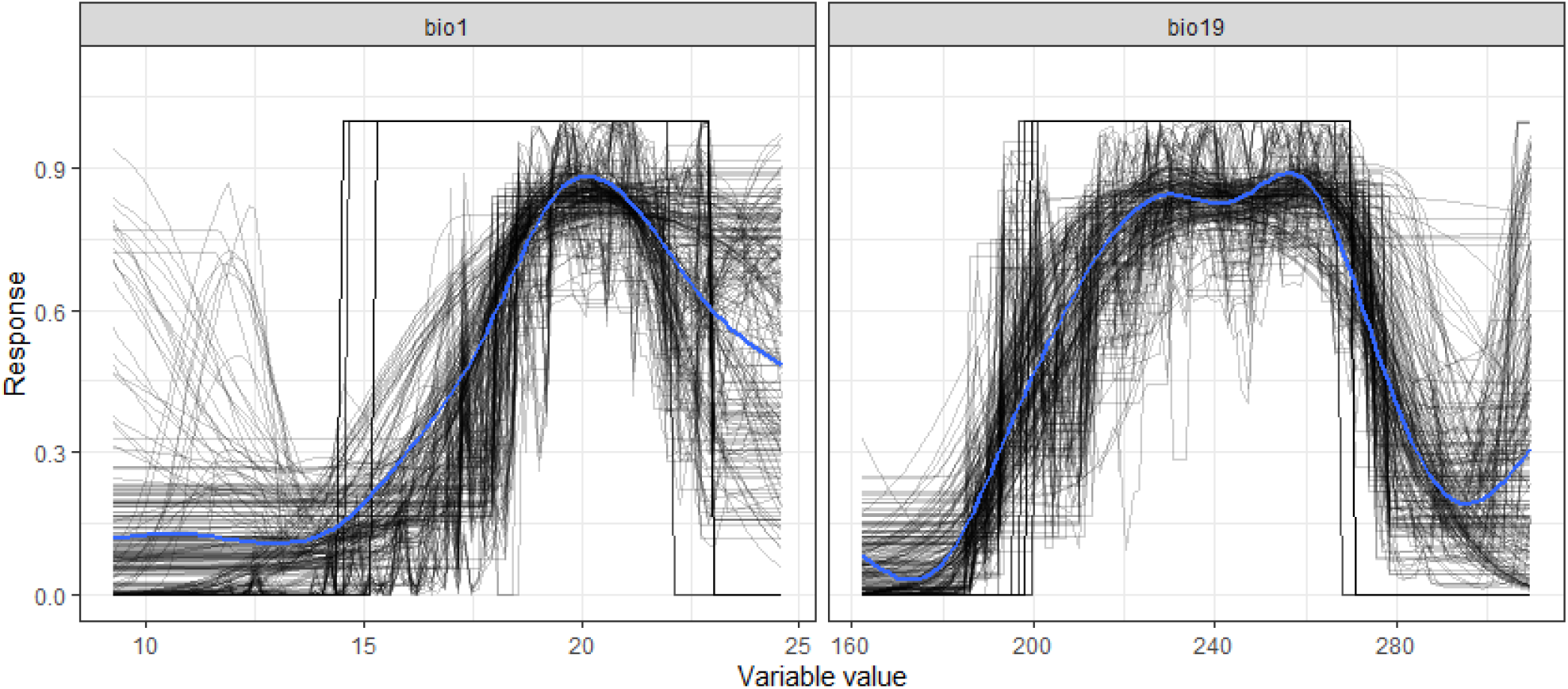
Response of *Phelsuma borbonica* to the selected Worldclim bioclimatic variables. The black lines represent the individual response curves for each iteration. The blue line represents the smoothed response across iterations. Bio1: annual mean temperature (°C); Bio19: precipitation of coldest quarter (mm).

**Fig. 2.**
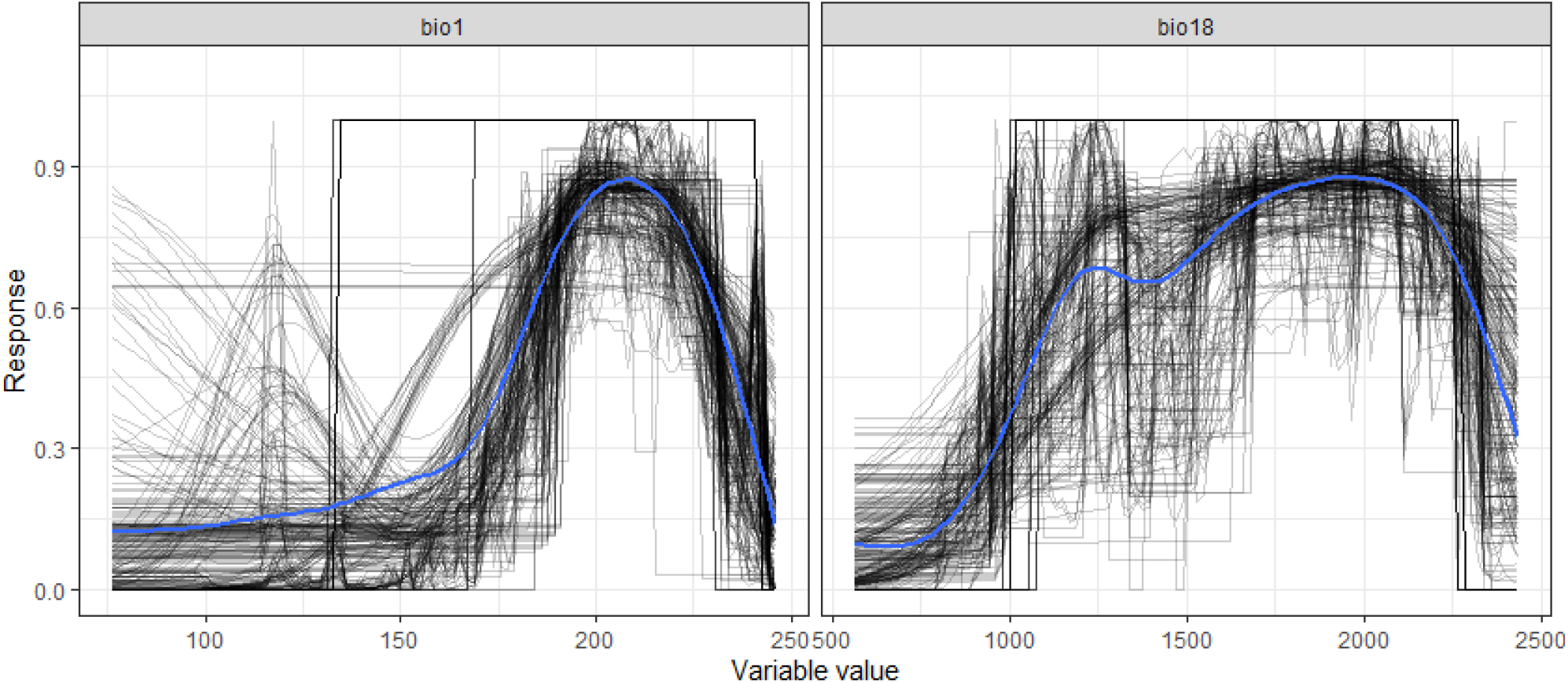
Response of *Phelsuma borbonica* to the selected CHELSA bioclimatic variables. The black lines represent the individual response curves for each iteration. The blue line represents the smoothed response across iterations. Bio1: annual mean temperature (°C ×10); Bio18: precipitation of warmest quarter (mm).

### Future distribution

Predictions differed strongly between modalities. Predictions derived from Worldclim baseline data indicated a dramatic decline in climate suitability across the entire island, regardless of the scenario or GCM (Fig. 3). The best predicted areas would shift upslope, but remain largely suboptimal. When based on CHELSA baseline data, predictions were more optimistic but highly variable between GCMs (Fig. 4). The ‘BCC-CSM-1-1’ GCM predicted little effect of climate change. The ‘MIROC5’ GCM indicated a slight decrease in climate suitability in the north-western part of the current distribution area, while the HadGEM2-AO predicted a strong decline in the eastern part.

**Fig. 3.**
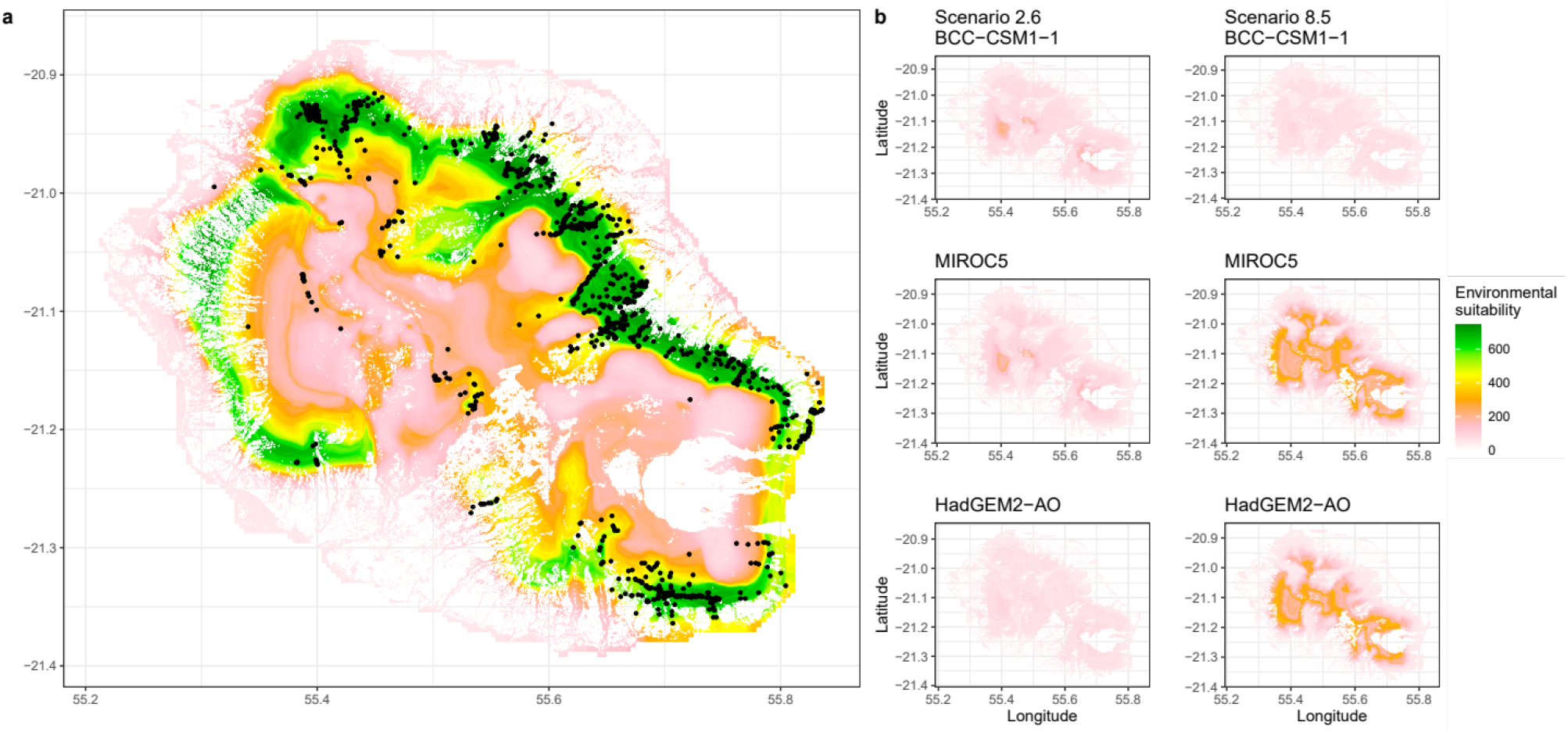
Current (a) and future (b) environmental suitability of *Phelsuma borbonica* in Reunion Island based on Worldclim baseline climate data.

**Fig. 4.**
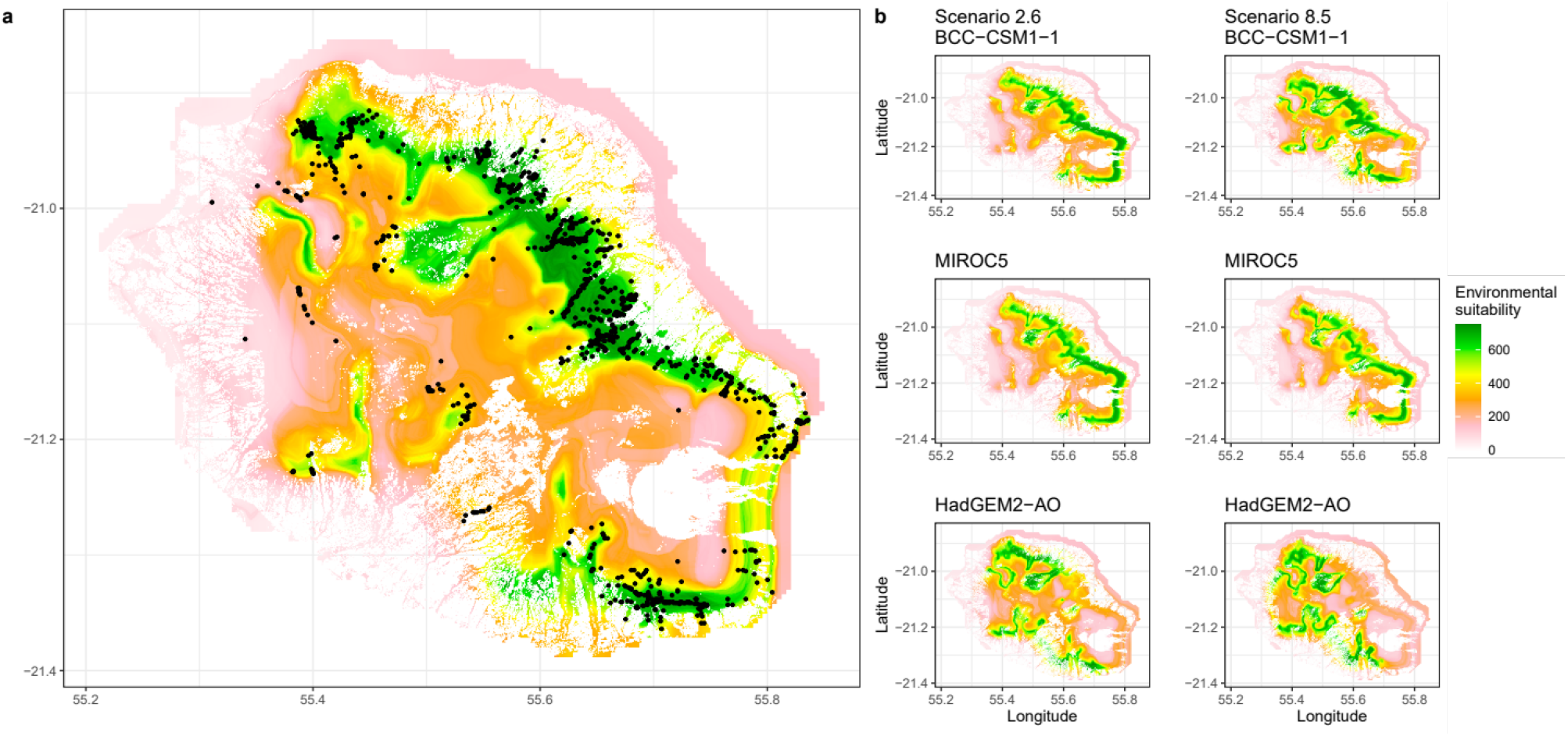
Current (a) and future (b) environmental suitability of *Phelsuma borbonica* in Reunion Island based on CHELSA baseline climate data.

### Effect of variable selection

After swapping the selected variables between baseline climate data, model performances decreased only slightly (median Boyce index (Worldclim) = 0.71; median Boyce index (CHELSA) = 0.70). The predicted current distribution was overestimated with Worldclim, and slightly underestimated with CHELSA. The differences between Worldclim and CHELSA in predictions of future climate suitability persisted. Predictions based on Worldclim strongly differed between GCMs (Fig. 5) but were consistent when based on CHELSA (Fig. 6).

**Fig. 5.**
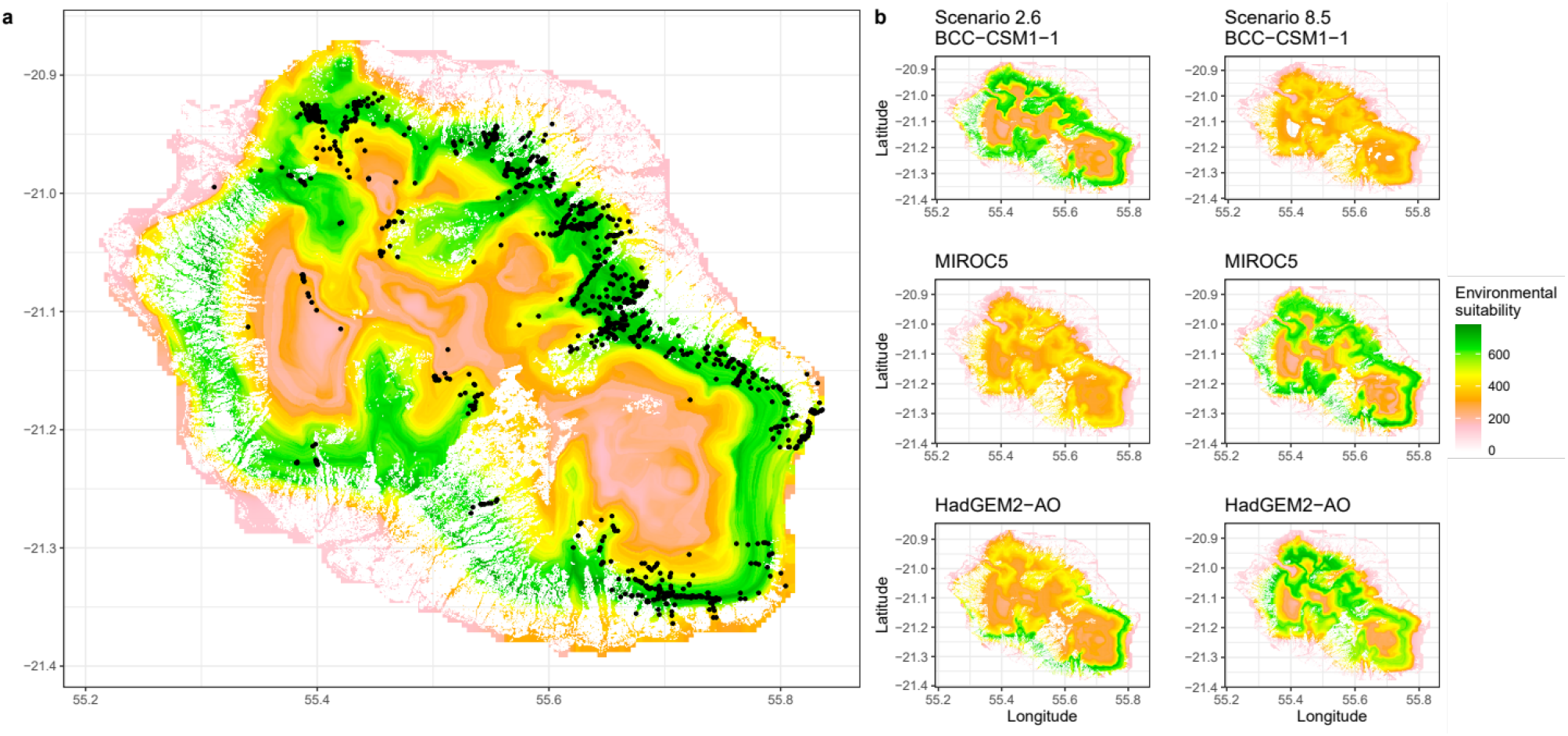
Current (a) and future (b) environmental suitability of *Phelsuma borbonica* in Reunion Island based on Worldclim baseline climate data using the selected variables from the CHELSA models.

**Fig. 6.**
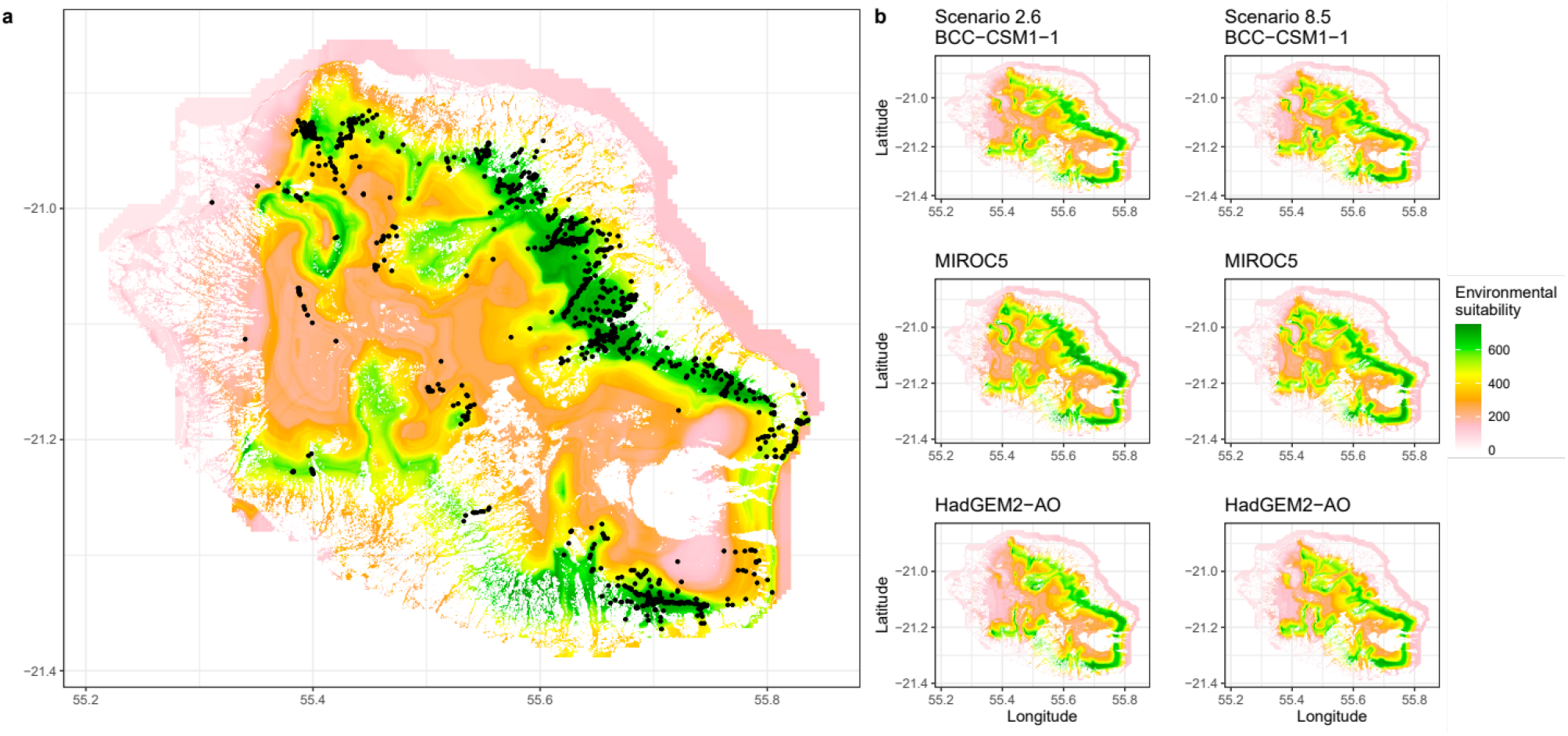
Current (a) and future (b) environmental suitability of *Phelsuma borbonica* in Reunion Island based on CHELSA baseline climate data using the selected variables from the Worldclim models.

### Uncertainty analysis

The highest source of uncertainty differed according to the metric of species range change considered (Fig. 7). Overall change in suitability scores were the most variable between the modelling technique (SDM), followed by the GCM and the baseline climate. Uncertainty related to sample bias correction and variable selection were of lower magnitude. Regarding the amount of spatial information shared between current and future predictions (overlap), the highest source of uncertainty was the modelling technique, followed by variable selection and baseline climate. The uncertainty related to the GCM was lower than that of baseline climate when considering overlaps. Sample bias correction was the lowest source of uncertainty.

**Fig. 7.**
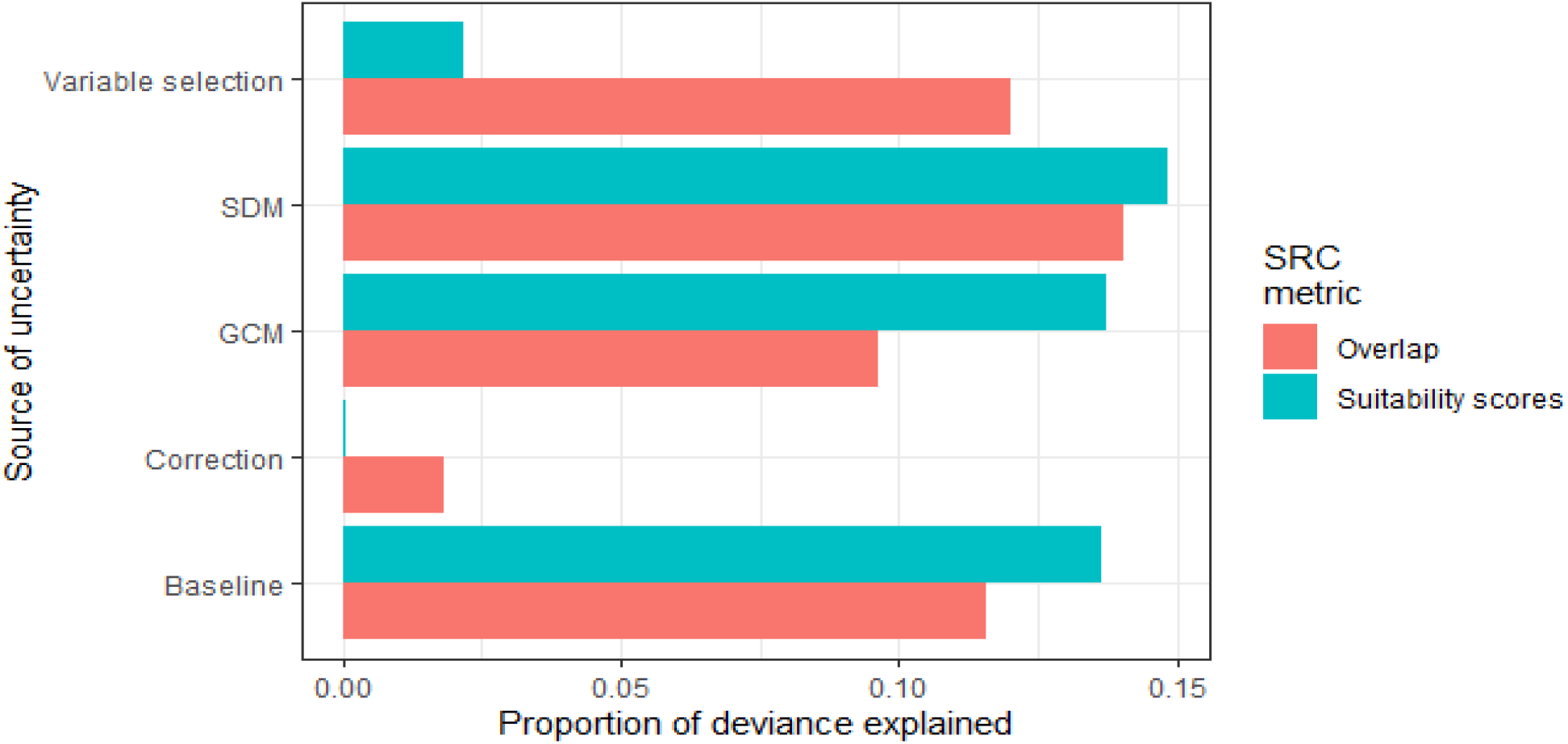
Proportion of deviance explained by five sources of uncertainty, quantified with two metrics of species range change (SRC). SDM: modelling technique (species distribution model); GCM: global circulation model; Correction: sample bias correction (non-random pseudo-absence selection); Baseline: baseline climate data (Worldclim versus CHELSA); Overlap: Schoener’s D overlap between current and future projections; Suitability scores: difference between summed suitability scores of current and future predictions.

The consensus maps indicate that the areas with the most consistent suitable conditions will be located at several places throughout the island (Fig. 8). The mean prediction maps also indicates that the best conditions will be largely suboptimal, from a maximum suitability score of 797 under current conditions to 362 by 2070.

**Fig. 8.**
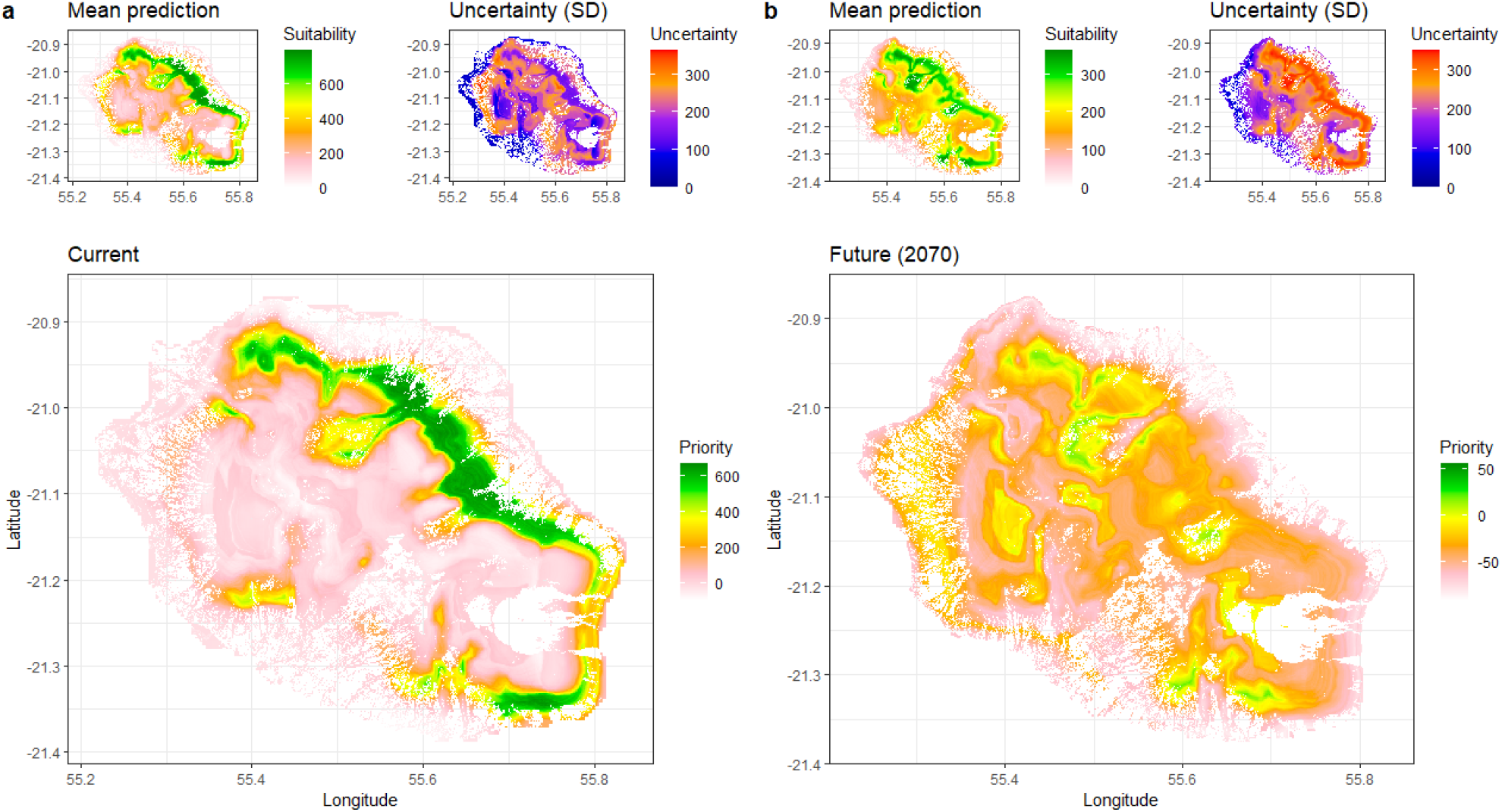
Consensus maps of current (a) and 2070 (b) priority areas for conservation for *Phelsuma borbonica*, derived from predicted climate suitability. Current and future predictions (bottom) are the result of mean projections (top left of each panel) discounted with inter-model variability (top right of each panel).

## Discussion

We predicted the future climate suitability of *P. borbonica* while accounting for multiple climate models and found a strong disagreement between future predictions derived from Worldclim and CHELSA baseline climate data. The uncertainty related to both methodological aspects and input climate data depended on the approach used to quantify species range change (summed suitability *versus* overlap). Mean projections indicate an important decline in climate suitability by 2070.

### Drivers of species distribution

The effect of temperature is consistent between simulations, with an optimal suitability estimated around 20°C on average throughout the year (mostly between 15 and 25°C, with a second peak near 10°C). The identified temperature window corresponds to the overall thermal tolerance of tropical reptiles (Sunday, Bates, & Dulvy, 2011).

*Phelsuma borbonica* is more present in areas with high precipitation, mostly corresponding to native remnant rainforests. We presume that *P. borbonica* benefits from a higher availability of specific native tree species (e.g., Arecaceae and Pandanaceae), which provide shelter, food and oviposition sites, as it is also found in other *Phelsuma* species from the Comoros and Seychelles (Noble et al., 2011; Augros et al., 2018; Augros, 2019). *P. borbonica* may also be adapted to cyclone activity, as this might be the case in Madagascar or in the Pacific islands (e.g., fallen trees used as hatching sites or open areas for thermoregulation; Ineich, 2010; Dubos et al., 2020). However, the species is mainly found in intact forests and may therefore use them as shelter to reduce cyclone impact instead.

Overall, *P. borbonica* can persist in very different habitats and environmental conditions, from cold and dry rocky slopes near the summits of the island (above 2.000 meters; Sanchez & Probst, 2017) to warm lowland and midland humid forests. We found a bimodality in the response curves to temperature and precipitation for several iterations. These were likely caused by the presence of isolated populations on mountain ridges and reflect the local adaptations to these specific environments. In Reunion, two subspecies were described, i.e. *P. borbonica borbonica* and *P. borbonica mater* (Meier, 1995; Probst & Deso, 2001). However, the two modalities we found do not correspond to these subspecies, which seem to share the same climatic niche instead. Preliminary unpublished genetic analyses suggest a strong isolation between these populations (Sanchez et al., 2015; unpubl. data). Further genetic analyses are needed to quantify the degree of isolation between populations.

Occurrence data which corresponded to isolated observations with no evidence of established populations (i.e., with no individual was observed after further surveys) were from areas that were predicted as unsuitable, a sign that our models identified well the species climatic niche.

Predictions based on Worldclim indicated that suitable conditions can be met around the island at intermediate altitudes. It is possible that *P. borbonica* once occupied this whole area, but was extirpated because of intensive agriculture and habitat fragmentation in the western part of the island. A local extinction of the *P. borbonica* was already documented in Cheke (1987). This depletion was associated with the severe deforestation that occurred in the 18th century at the intermediate and lower belt of the island, and a high pressure from invasive alien species (i.e., the introduced wolf-snake *Lycodon aulicus;* Cheke & Hume, 2010). These combined factors may have dramatically decreased habitat suitability and habitat availability for *P. borbonica* in the past centuries, which may explain its absence in some of the predicted climatically suitable areas. The former presence of *P. borbonica* in the western part of the island is supported by the presence of isolated remnant populations in the southwest (near Les Makes & Le Tampon) and recently extinct populations in the northern lowlands (Cheke, 1987).

As shown by the response curves, *P. borbonica* persists over a relatively wide range of climatic conditions. However, its presence remains localised throughout the island. To fully understand and explain the current distribution patterns of *P. borbonica*, there is a need to also consider additional key factors such as microhabitat use and behaviour (Kearney, Shine, & Porter, 2009; Porter & Kearney, 2009). For instance, coldblooded species can respond to climate change by altering their period of activity and thermoregulation time (Kearney, Shine, & Porter, 2009). Moreover, the availability of suitable oviposition sites, thermoregulation sites with specific exposure to wind, sun or rain is a strong determinant of gecko occupancy (Ineich, 2010; Bungard et al., 2014; Augros et al., 2017, 2018). Habitat structure is provided in native forests by tree species from the Arecaceae and Pandanaceae families and abiotic features (e.g. sunny-exposed rocks, cliffs, manmade structures; Petren & Case, 1998; Augros et al., 2017). In additions to climate change, habitat modifications, such as further urbanisation and deforestation will strongly influence the future distribution of *P. borbonica* in Reunion Island.

Biotic interactions play an important role in shaping species distributions (Araújo et al., 2007). The current distribution of *P. borbonica* is strongly influenced by the occurrence of invasive alien species. The wolf snake *Lycodon aulicus* is presumed to be the cause of historical local extinctions through predation (Cheke, 1987). The introduced Giant Madagascar day gecko *P. grandis* and the Gold-dust day gecko *P. laticauda* are present throughout the island. Both species were reported from areas within the distribution of *P. borbonica*. To date, there is no evidence of local extirpation of *P. borbonica* by these two species. However, *P. grandis* was suspected to be the cause of local extinctions of *Phelsuma* species in Mauritius, presumably through competitive exclusion and/or predation (Buckland et al., 2014). It is also the case for *P. laticauda* in French Polynesia (Lund, 2015). *Phelsuma laticauda* is a rising cause of concern in the south of Reunion Island where it possibly threatens the persistence of the critically endangered *P. inexpectata* (NOI, unpubl. data; but see Porcel et al., 2021). Other introduced species may also affect *P. borbonica* through predation and/or habitat alterations, including other reptiles (*Agama agama, Calotes versicolor, Furcifer pardalis*), rodents (*Rattus rattus, Mus musculus*), ants (*Solenopsis geminata*), birds (*Pycnonotus jocosus, Acridotheres tristis*) and plants (*Lantana camara*). The rate of invasion is increasing in Reunion Island (Fenouillas et al., 2021). Further studies will need to account for biotic interactions to better understand the key drivers of the distribution of *P. borbonica* and refine forecasted predictions.

### Sources of uncertainty

We found a strong uncertainty driven by the modelling technique and GCMs. This is consistent with the findings of Buisson et al. (2010) and Baker et al. (2016) and advocates the use of a wide range of modelling techniques and GCMs for conservation planning. However, the uncertainty related to the baseline climate data was stronger than that of GCM when considering the spatial overlap between current and future predictions. When considering overall suitability, the importance of the baseline climate was probably underestimated because it affected the suitability scores in a lower extent than GCMs, however, resulted in higher discrepancies in spatial distribution of the future suitable conditions. The mismatch caused by the baseline climate data can be due to the differences in temporal coverage, with Worldclim representing the conditions of the 1960-1990 period while CHELSA was computed for 1979-2013. However, temporal coverage cannot fully explain these discrepancies because future predictions (both for 2070) also strongly differed, even when using the same predictors. Another factor potentially affecting these differences in predictions is the higher accuracy in predictions for precipitation of the CHELSA climatic data, with more topographic factors accounted for (Karger et al., 2017). This difference in accuracy may be exacerbated in mountainous environments such as Reunion Island. These results suggest that the spatial component of species range change should not be neglected when the aim is to identify priority areas for conservation. More generally, the importance of drivers of uncertainty may be downplayed when ignoring spatial information. We recommend the use of multiple approaches, including overlap estimations, in uncertainty assessments of species range change.

### Conservation considerations

The current distribution of *P. borbonica* is generally consistent with the predicted future suitable areas (see Fig8b, ‘Mean prediction’ panel). We recommend that the areas identified with the highest certainty should be prioritised for conservation and habitat restoration. In the context of Reunion Island, conservation actions are drastically limited by land use policy as the available land is strongly disputed for by urbanisation and agriculture planning. Overall, conservation efforts should be intensified in the forested uplands, where predictions are favourable and anthropogenic pressure is the least. Nevertheless, the conservation of the small, isolated populations of *P. borbonica* along some of the mountain ridges and in the west of Reunion Island is of paramount importance, as these represent remnant populations that possibly form genetically isolated entities (Sanchez et al., 2015; unpubl. data). Being supposedly more adapted to colder conditions, mountain ridge populations may also be at even greater risk due to climate change (Raxworthy et al., 2008; Freeman et al., 2018). We recommend the close monitoring of these populations for early detection of potential signs of population declines.

Conservation actions should encompass a range of management strategies, including the protection of native forests, restoration of degraded habitats, creation of artificial oviposition sites and the implementation of sustainable agricultural practices. The control of invasive species represents an additional challenge, for instance with *P. grandis* and *P. laticauda* currently in expansion throughout the island (Dubos, 2013; Porcel et al., 2021).

This study stresses the need for proactive conservation actions given the high risk of extinction predicted by some of our models. *Phelsuma borbonica* is already threatened by habitat loss and fragmentation, which will likely increase in the future with human population. Future conservation actions will need to consider socioeconomic factors to prevent potential land use conflicts (Lagabrielle et al., 2011). This can be achieved by involving stakeholders from urban and agricultural sectors and conservation practitioners into public decision-making processes.

Depending on the climate data considered, our models predicted either a strong decline throughout the entire island, upward, westward, or eastward shifts, or almost no change. Despite the high uncertainty, we identified the areas with the most consistent predictions of suitable climate by 2070. In a context of urgent decision making, we advocate the use of all the available tools to prevent possible extinctions in spite of the apparent uncertainty. Forecasting models need to consider a wide range of methods and data, and assess the variability between them in order to identify and mitigate potential sources of uncertainty and provide relevant conservation guidelines.

## Aknowledgements

We thank Boris Leroy for useful discussions. We are thankful to all the field workers that collected the data used in this study. We thank Biotope Océan Indien and Cynorkis for providing additional data.

## Data accessibility statement

All the occurrence data and the R scripts used in this study are available in online supporting information.

## Notes

### Competing Interest Statement

The authors have declared no competing interest.

